# Heterotypic droplet formation by pro-inflammatory S100A9 and neurodegenerative disease-related alpha-synuclein

**DOI:** 10.1101/2024.11.27.625672

**Authors:** Dominykas Veiveris, Aurimas Kopustas, Darius Sulskis, Kamile Mikalauskaite, Marijonas Tutkus, Vytautas Smirnovas, Mantas Ziaunys

## Abstract

Liquid-liquid phase separation (LLPS) of proteins and nucleic acids is a rapidly emerging field of study, aimed at understanding the process of biomolecular condensate formation and its role in cellular functions. LLPS has been shown to be responsible for the generation of promyelocytic leukemia protein bodies, stress granules, and intrinsically disordered protein condensates. Recently, it has been discovered that different neurodegenerative disease-related proteins, such as alpha-synuclein (related to Parkinson’s disease) and amyloid-beta (Alzheimer’s disease) are capable of forming heterotypic droplets. Other reports have also shown non-LLPS cross-interactions between various amyloidogenic proteins and the resulting influence on their amyloid fibril formation. This includes the new discovery of pro-inflammatory S100A9 affecting the aggregation of both amyloid-beta, as well as alpha-synuclein. Combined, these observations suggest that protein interactions during LLPS and heterotypic droplet formation may be a critical step in the onset of neurodegenerative diseases.

In this study, we explore the formation of heterotypic droplets by S100A9 and alpha-synuclein using a range of different spectroscopic and microscopic techniques. We show that the protein mixture is capable of assembling into both homotypic, as well as heterotypic condensates and that this cross-interaction alters the aggregation mechanism of alpha-synuclein. In addition, it also stabilizes a specific fibril conformation, which has a higher propensity for self-replication. These results provide insight into the influence of S100A9 on the process of neurodegenerative disease-related protein LLPS and aggregation, bringing us one step closer to developing a potential cure or treatment modality.

## Introduction

Protein and nucleic acid liquid-liquid phase separation (LLPS) is a process during which biomolecules condense into high concentration membraneless droplets^1,2^. This phenomenon has recently gained recognition due to its role in various biological processes, including transcription regulation, genome organization and immune response ^1,3–5^. However, recent studies have also shown that aberrant LLPS might be associated with the onset of neurodegenerative disorders, such as Alzheimer’s ^6^ or Parkinson’s disease ^7^, as well as various forms of cancer ^3,4,8,9^. Despite enormous progress in this field, the mechanism and implications of LLPS are still far from being fully understood, with new insight being discovered on a regular basis ^10–12^. Due to its function in not only the regulation of countless biological processes, but also manifestation of several widespread diseases, it is imperative to gain a deeper insight into biomolecule condensate formation.

Over the last few years, it has been discovered that a number of different proteins can assemble into heterotypic droplets, i.e. condensates composed of two structurally distinct molecules ^13–15^. In the case of neurodegenerative disorders, this cross-interaction has been hypothesized as a possible intermediate step in the onset of amyloid diseases ^13^. Several amyloidogenic protein pairings were observed to form heterotypic condensates, including alpha-synuclein (α-syn) with Tau ^16,17^ and TDP-43 ^18^, as well as prion proteins with Tau ^19^ and α-syn ^20^. The amyloid-beta peptide ^21^ has also been shown to associate into heterotypic droplets with proteins containing low complexity domains ^22^. Most of these protein pairings were reported to cross-interact and affect each other’s aggregation under non-LLPS conditions as well ^15,23–28^. Combined with studies describing amyloid plaques in disease-affected brains having a heterogenous protein content ^29–31^, it is possible that heterotypic droplet formation plays a critical role in the onset and progression of several neurodegenerative disorders.

In recent years, it has also been discovered that there exists a cross-interaction between α-syn and S100A9 ^32^. α-syn is an intrinsically disordered presynaptic protein, whose aggregation into Lewy bodies and Lewy neurites is associated with the second most prevalent neurodegenerative disorder – Parkinson’s disease ^33–35^. α-syn has been the subject of numerous LLPS studies because of its ability to readily form protein droplets in vitro under high molecular crowding conditions ^7,36,37^. S100A9 is part of a calcium-binding pro-inflammatory S100 protein family ^38^. Due to the protein’s ability to interact with amyloid-beta, α-syn and Tau, as well as its own amyloidogenic properties ^32,39,40^, it is considered that S100A9 may play a critical role in the onset of several neurodegenerative disorders. Recent studies have shown that S100A9 can significantly alter the aggregation pathway of α-syn, leading to the stabilization of a specific fibril secondary structure ^32,41^. In contrast to α-syn, there are currently no reported data on whether S100 family members can undergo LLPS to a comparable extent as other amyloidogenic proteins.

For this reason, our study was dedicated to examining the cross-interaction between α-syn and S100A9 in the context of protein condensate formation. In this work, we demonstrate the ability of α-syn and S100A9 to form both homotypic and heterotypic droplets under high molecular crowding conditions. This cross-interaction influences the aggregation kinetics of α-syn and stabilizes a single fibril conformation. In addition, the resulting strain of fibrils has a notably higher self-replication propensity, when compared to aggregates formed in homotypic α-syn droplets. Combined, these results suggest that heterotypic condensate formation by the pro-inflammatory S100A9 and α-syn is not only possible, but may also be an important factor in the onset of neurodegenerative disorders.

## Materials and Methods

### Cloning

The mCherry and S100A9 genes were amplified and fused using standard PCR methods. The products were inserted into a pET28a(−) vector via NcoI and BamHI restriction sites by standard cloning techniques ^42^ yielding mCherry-S100A9 construct with (GGGGS)_2_ linker between the genes and N-terminal (His)_6_ tag. Primers used in this study can be found in Table S1.

### Protein purification

Recombinant α-syn was purified as described previously ^43^. During the last purification step with size-exclusion chromatography (SEC), the protein was exchanged into PBS (pH 7.4) and stored at - 20°C. After all SEC cycles were completed, the protein fractions were thawed at 4°C, combined and concentrated to 600 µM using 10 kDa protein concentrators. The prepared protein solutions were then divided into 0.5 mL aliquots and stored at -20°C prior to use. All experimental procedures in this work were carried out by using the same batch of α-syn. The eGFP-labeled α-syn was purified identically, with the exception of using 70% saturation ammonium sulphate in the protein precipitation step ^44^.

S100A9 was purified as described previously ^45^. After gel filtration in PBS buffer (pH 7.4), the protein was concentrated to 500 µM, aliquoted and stored at -80°C prior to use. mCherry-S100A9 was purified according to the S100A9 protocol with immobilized metal affinity replacing anion exchange chromatography ^41^.

### Liquid-liquid phase separation

Polyethylene glycol (PEG, 20 kDa average molecular weight) was combined with MilliQ H_2_O and 10x PBS to a final concentration of 40% (w/v) and 1x PBS. pH adjustments to 7.4 were done by adding a concentrated sodium hydroxide solution. Due to the high viscosity of the solution, it was vigorously mixed with magnetic stirring (900 RPM) during the pH measurement procedure. The solution was filtered through a 0.22 µm pore-size syringe filter and stored at 4°C prior to use.

To induce protein LLPS, the α-syn and S100A9 solutions were combined with 1x PBS (pH 7.4), 40% PEG (pH 7.4) and fluorescently labeled protein stock solutions. The final reaction mixtures contained 20% PEG, 1% labeled protein (either 2 µM eGFP-α-syn or 2 µM mCherry-S100A9), 200 µM α-syn and 0, 50 µM S100A9 concentrations. Control solutions were prepared without the addition of either α-syn or S100A9. Due to the importance of the component mixing order ^46^, the PBS and PEG solutions were combined first, after which the proteins were added (α-syn first, S100A9 second, labeled proteins third).

### Fluorescence and brightfield microscopy

15 μL aliquots of each sample were pipetted onto 1 mm-thick glass slides (Fisher Scientific, cat. No. 11572203), covered with 0.18 mm coverslips (Fisher Scientific, cat. No. 17244914) and imaged as described previously ^44^ using an Olympus IX83 microscope with a 40x objective (Olympus LUCPLANFL N 40x Long Working Distance Objective) and fluorescence filter cubes (470 – 495 nm excitation and 510 – 550 nm emission filters for eGFP-α-syn, 540 - 550 nm excitation and 575 - 625 nm emission filters for mCherry-S100A9). For fluorescence microscopy images, identical background subtraction and contrast/brightness settings were applied to all images. For brightfield microscopy images, only the contrast/brightness settings were adjusted. Data analysis was done using ImageJ software ^47^.

To examine samples containing both fluorescently labeled proteins using two-color fluorescence microscopy, the solutions were placed on cleaned glass coverslips (Menzel Coverslip 24×60mm #1.5 (0.16 – 0.19 mm), Thermo Scientific, cat. no. 17244914). For this, a 100 µL droplet suspension containing 200 µM α-syn and 1% of eGFP-α-syn, mCherry-S100A9 or both was slowly added on a bare glass surface using wide-orifice tips (Finntip 250 Wide, Thermo Scientific, cat. no. 9405020) with no pipetting and any subsequent washing of the sample. The miEye, a home-built super-resolution imaging system ^48^, was employed to visualize these fluorescent samples. All experiments were conducted in a TIRF imaging mode with a quad line beamsplitter R405/488/561/635 (F73-866S, AHF Analysentechnik) mounted in the microscope’s body. 488 nm and 561 nm lasers (Integrated Optics) were used to excite the fluorescently tagged α-syn droplets attached to the glass surface. The emission pathway of miEye was modified to a dual-view regime by inserting a 550 nm long-pass dichroic mirror into the Fourier space present in the microscope’s 4f configuration part. This resulted in the two spectrally distinct emission light collecting channels which, for simplicity, here we refer them to as eGFP channel and mCherry channel. The eGFP channel was equipped with a 525/45 band-pass filter, whereas the mCherry one – with a 697/75 band-pass filter. Both channels were projected and imaged on a single industrial CMOS camera (Alvium 1800 C-511m, Allied Vision Technologies) with its exposure time set to 100 ms. Data analysis was done using ImageJ software ^47^, example is shown as Figure S1.

### Droplet statistical analysis

For each condition, a total of thirty 500×500 pixel size images (1 pixel – 325 nm) were obtained (available at: https://data.mendeley.com/datasets/tvf9nwtdhn/1). The statistical analysis was done as described previously ^46^. In brief, droplet parameters were analyzed with ImageJ software ^47^ using automatic threshold selection and particle analysis. All particles of 4 or less pixel size were regarded as artefacts and not taken into account. The total droplet count was the sum of all particles detected in all 30 images for each condition. Average droplet volume was calculated based on the particle areas (assuming completely spherical condensates). Data analysis was done using Origin software and an ANOVA One-way Bonferroni means comparison (n=30).

### S100A9 fibril preparation

S100A9 fibrils were prepared using a previously described protocol ^49^, which generates worm-like amyloid aggregates^50^. The protein stock solution was diluted to 200 µM using 1x PBS (pH 7.4). The reaction solution was then placed in a 2.0 mL non-binding test tube (1 mL solution) and incubated under quiescent conditions at 37°C for 24 hours. The prepared aggregate solution was then stored at 4°C. Before further experimental procedures, the fibril solution was concentrated to 400 µM using 0.5 mL volume 10 kDa Pierce protein concentrators.

### LLPS and aggregation kinetics

Solutions containing 20% PEG (w/v), 100 µM thioflavin-T (ThT), 200 µM α-syn and 0, 5 or 50 µM of non-aggregated or fibrillar S100A9 were distributed to 96-well non-binding plates (100 µL volume in each well, 4 repeats for every condition), sealed with Nunc sealing-tape and incubated under quiescent conditions at 37°C in a ClarioStar Plus plate reader. Fluorescence intensity measurements were performed every 10 minutes. ThT fluorescence intensity was monitored using 440 nm excitation and 480 nm emission wavelengths. Due to the time required for sample preparation and distribution procedures, the first measurement was performed approximately 30 minutes after the mixtures were prepared. During this time, the samples were kept at room temperature (22°C). All data analysis was done using Origin software.

### Aggregate reseeding

After the initial LLPS and aggregation reactions, the solutions from each of the 4 repeats were combined and centrifuged at 12 000 x g for 20 minutes. The supernatants were then carefully removed and replaced with an identical volume of PBS (7.4). The centrifugation and resuspension procedure was repeated three times in order to separate the aggregates from the initial reaction solutions. For the reseeding reactions, the α-syn stock solution was combined with PBS (pH 7.4), ThT and the resuspended aggregates to final reaction mixtures containing 200 µM α-syn, 100 µM ThT and 10% (v/v) aggregate solutions. The reactions were monitored as described previously under quiescent conditions and 37°C. After 24 hours, the samples from each of the 4 repeats were combined and the entire reseeding procedure was repeated for a second time. The final resulting samples were then used for electron microscopy.

### Cryo electron microscopy (Cryo-EM)

For cryo-EM sample preparation, 3 μL of alpha synuclein fibrils were applied to the glow-discharged holey carbon Cu grids (Quantifoil) and blotted with filter paper using Vitrobot Mark IV (FEI Company). The grids were immediately plunge-frozen in liquid ethane and clipped. Cryo-EM data was collected on Glacios transmission electron microscope (Fisher Scientific) operated at 200 kV and equipped with a Falcon IIIEC camera. The micrographs were aligned, motion corrected using MotionCorr2 1.2.1 ^51^ and the contrast transfer function was estimated by CTFFIND4 ^52^. The fibrils were picked and all subsequent 2D classifications were performed in Relion 5.0 ^53^. Distribution of polymorphs was identified by FilamentTools (https://github.com/dbli2000/FilamentTools) as a part of Relion software. Cryo-EM data collection and 2D classification statistics can be found in Table S2.

## Results

### Heterotypic droplet formation

Previous reports of S100A9 and α-syn cross-interactions ^32,41^, as well as the ability of α-syn to form heterotypic condensates ^16–18,20^, has prompted the need to examine this specific protein pairing in the context of liquid-liquid phase separation. In order to test the hypothesis of heterotypic droplet formation, high concentration α-syn samples were combined with a 100-fold lower concentration of either eGFP-α-syn (control) or mCherry-S100A9. To enhance the level of condensate formation, the protein solutions were supplemented with 20% (w/v) of a molecular crowding agent – polyethylene glycol (PEG, 20 kDa) ^44^. The samples were then imaged using fluorescence microscopy and a total of 30 images for each were obtained and used for particle count and volume distribution analysis. If the hypothesis is correct, both eGFP-α-syn, as well as mCherry-S100A9 should be incorporated into the α-syn droplets and the fluorescence images would show a similar distribution of protein condensates. Oppositely, if mCherry-S100A9 could not interact with α-syn, we would observe either no droplet formation or condensates assembled from only the labeled protein.

When the samples were analyzed, both α-syn with GFP-α-syn (Figure 1A) and α-syn with mCherry-S100A9 (Figure 1B) solutions contained a large number of droplets with varying size. Surprisingly, analysis of the images revealed that the sample containing mCherry-S100A9 had a significantly higher number of particles (Figure 1F, n=30, p<0.001). However, the average particle volume was not significantly different (Figure S2A, n=30, p<0.001), despite having a lower mean (∼15 µm^3^ as opposed to ∼22 µm^3^). It is worth noting that the determined apparent particle volume may be influenced by the image acquisition technique and subsequent image processing. To determine if this peculiar effect on the condensate number is not related to the self-assembly of the fluorescently labeled proteins, an identical analysis was conducted on samples containing only 2 µM of eGFP-α-syn or mCherry-S100A9 (Figure 1C, D).

**Figure 1.**
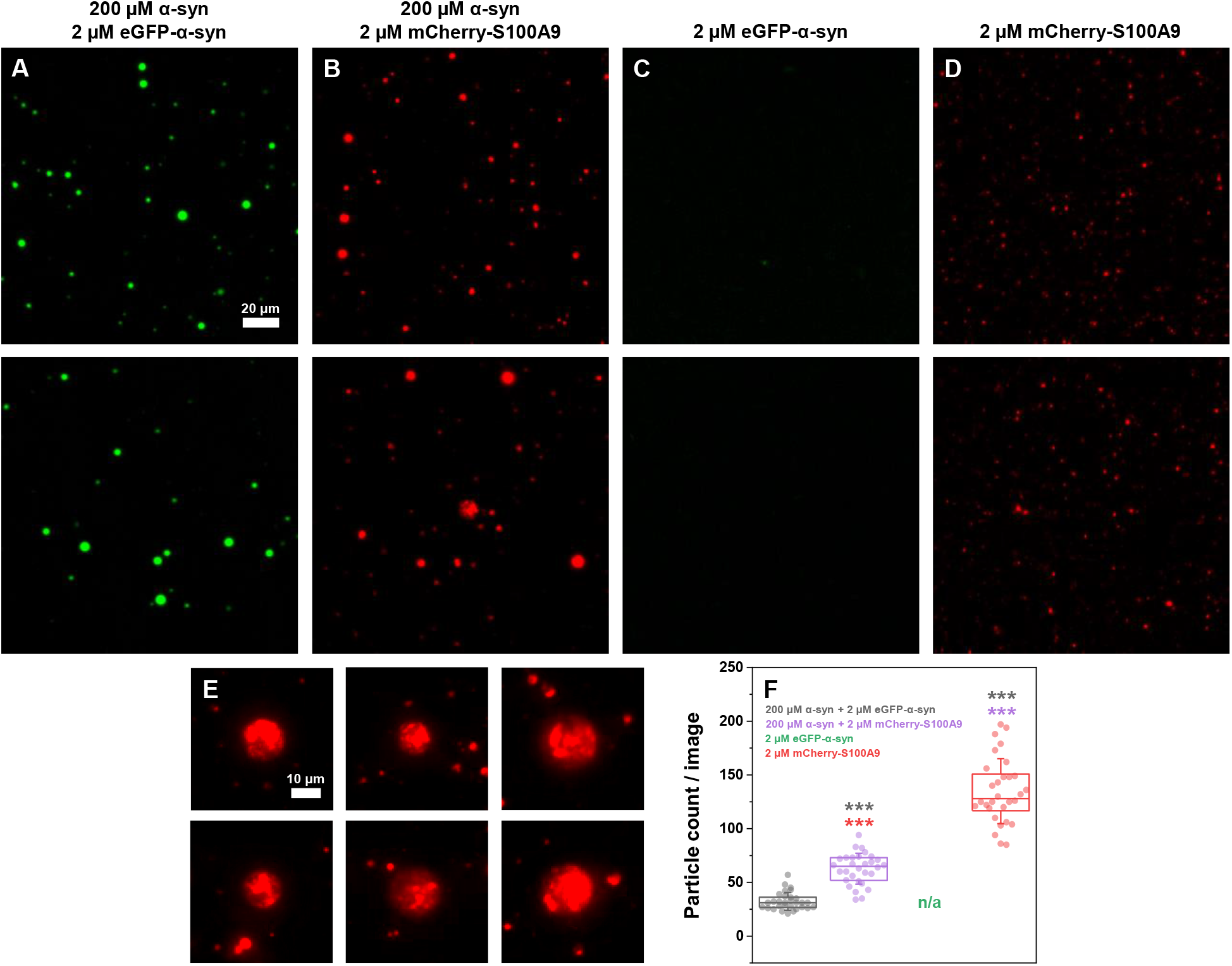
Fluorescence microscopy images of alpha-synuclein (α-syn) and labeled protein condensate formation. Two representative images of 200 µM α-syn with either 2 µM eGFP-α-syn (A) or 2 µM mCherry-S100A9 (B, scale bar – 20 µm). Two representative control sample images of 2 µM eGFP-α-syn (C) and 2 µM mCherry-S100A9 (D, scale bar – 20 µm). Images of unevenly filled droplets from 200 µM α-syn with 2 µM mCherry-S100A9 images (E, scale bar – 10 µm). Statistical analysis (ANOVA Bonferroni means comparison, thirty 500×500 pixel size images, ns-not significant, ***-p<0.001) of the particle count per image (F). The 2 µM eGFP-α-syn images did not contain a sufficient number of particles for statistical analysis. Additional fluorescence microscopy images (Olympus IX83 microscope) are available as online material.

As expected, eGFP-α-syn did not form any notable number of assemblies, however, the mCherry-S100A9 sample images contained a large quantity of small particles. Data analysis revealed that the mCherry-S100A9 sample was comprised of a significantly larger number of particles than both α-syn with eGFP-α-syn, as well as α-syn with mCherry-S100A9 samples (n=30, p<0.001). The average particle volume was also significantly lower than in both other samples (Figure S2A). These findings indicate that the fluorescently labeled S100A9 forms visible/detectable particles even at a low concentration. Their self-association into these small particles, along with their incorporation into heterotypic droplets could account for the higher number of condensates detected in the α-syn + mCherry-S100A9 sample.

Another interesting phenomenon observed in the α-syn with mCherry-S100A9 sample was the formation of unevenly filled heterotypic droplets (Figure 1E). Upon closer inspection, while the droplets had a faintly visible spherical shape, the fluorescently labeled S100A9 was not evenly distributed within them, forming areas of lower and higher fluorescence intensity. In contrast, the larger eGFP-α-syn sample droplets all displayed an even fill of the fluorescently labeled protein (Figure S3). These results suggested that, despite the cross-interaction of both proteins and the formation of heterotypic condensates, S100A9 still retained a higher tendency to self-associate even within the droplets.

In order to determine if the protein condensates were all heterotypic, or if the droplets could also be homotypic, samples containing α-syn with both fluorescently labeled proteins were examined using two-color fluorescence microscopy. Overlaid two-color images revealed perfect co-localisation, and varying levels of either protein (Figure 2A-C). The majority of small particles were composed mainly of mCherry-S100A9, while generally larger droplets contained either only eGFP-α-syn or both labeled proteins.

**Figure 2.**
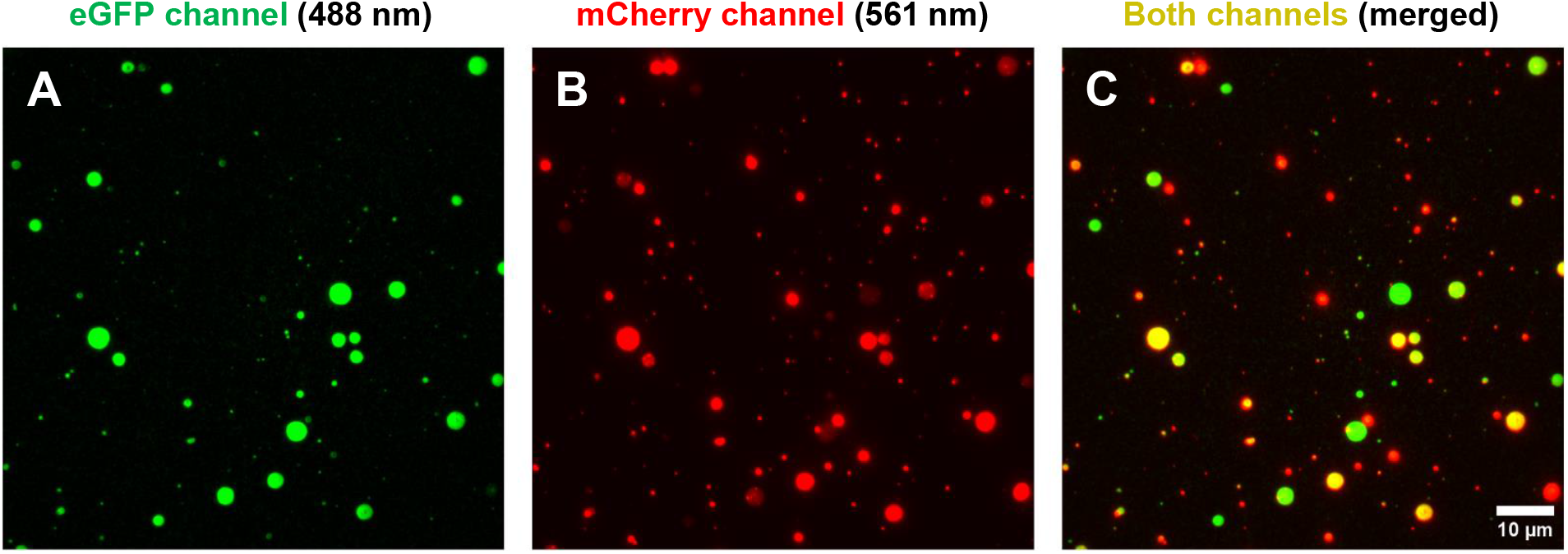
TIRF microscopy images of the surface-immobilized α-syn and α-syn-eGFP or mCherry-S100A9 droplets. Using a 488 nm laser illumination (miEye microscope), the fluorescence of eGFP-α-syn and mCherry-S100A9 was observed in the eGFP channel (A), whilst exciting this sample with a 561 nm laser yielded the fluorescence visible in the mCherry channel (B). These two images were acquired on the same surface position of the sample. Overlaid eGFP and mCherry channel images (C) show the colocalization of such droplets at some surface locations.

Since both proteins displayed the ability to form heterotypic droplets, it was further examined how a larger concentration of unlabeled S100A9 would influence the process of α-syn LLPS. Based on the results, the first notable distinction was that the sample with both proteins (200 µM α-syn and 50 µM S100A9) appeared to contain a larger number of particles (Figure 3A, B). Data analysis of each sample’s images confirmed this observation (Figure 3D). The sample average volume distribution followed a similar trend as in the case of α-syn with mCherry-S100A9 (Figure S2B), where the sample with both proteins had a lower, yet not significantly different mean value.

**Figure 3.**
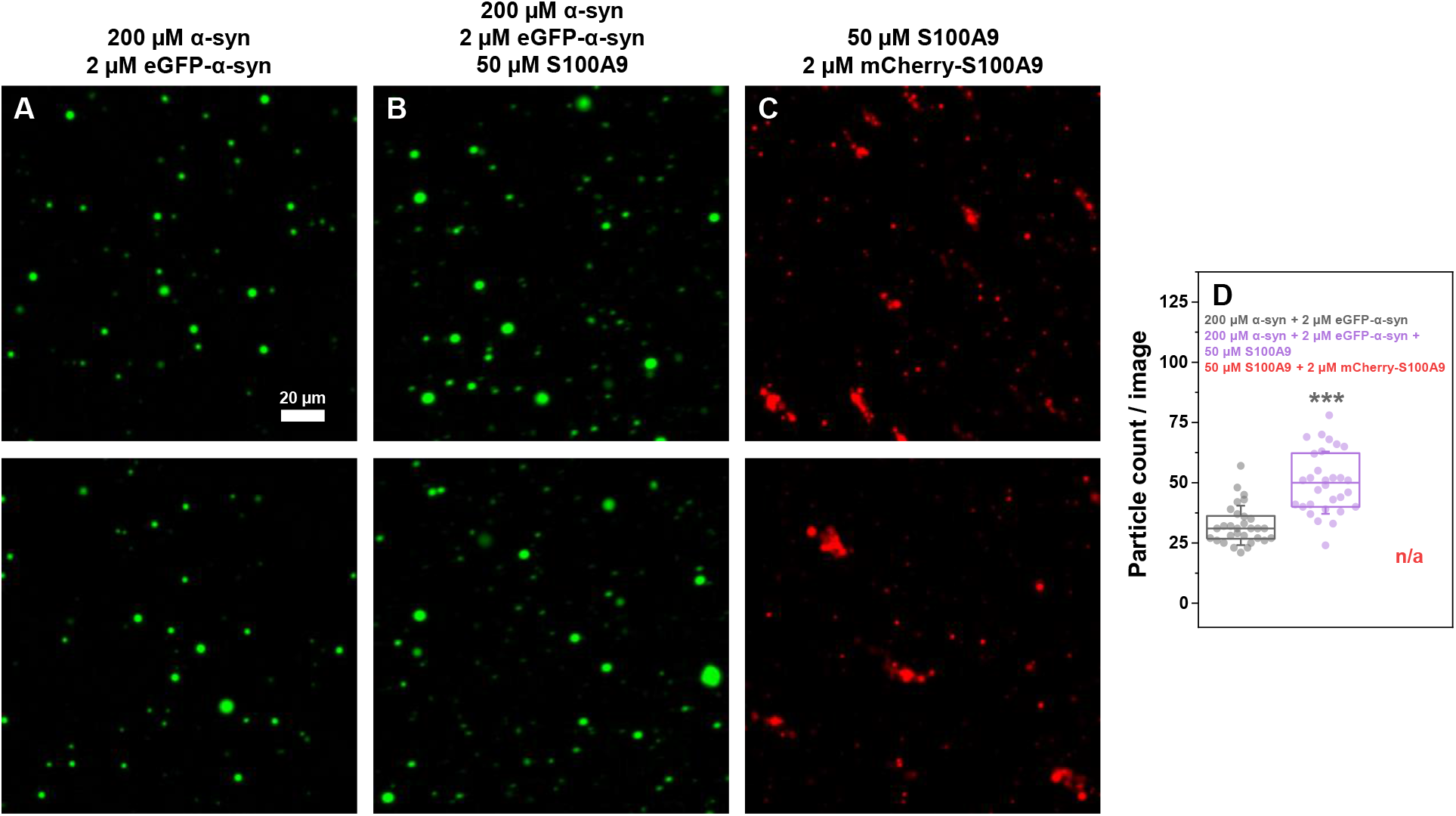
Fluorescence microscopy images of alpha-synuclein (α-syn) and S100A9 condensate formation. Images of 200 µM α-syn with 2 µM eGFP-α-syn (A), 200 µM α-syn with 2 µM eGFP-α-syn and 50 µM S100A9 (B) or 50 µM S100A9 with 2 µM mCherry-S100A9 (C, scale bar – 20 µm). Statistical analysis (ANOVA Bonferroni means comparison, thirty 500×500 pixel size images, ns – not significant, *** -p<0.001) of the particle count per image (D). Box plots indicate the interquartile range, error bars are for one standard deviation (n=30). The 50 µM S100A9 with 2 µM mCherry-S100A9 images contained both droplets and aggregates, which prevented an accurate statistical analysis. Additional fluorescence microscopy images (Olympus IX83 microscope) are available as online material.

Interestingly, when the sample did not contain α-syn, S100A9 with mCherry-S100A9 formed a mixture of droplets and various amorphous aggregates (Figure 3C). The presence of these structures prevented an accurate statistical analysis and also raised questions regarding the nature of the cross-interaction between both proteins. Taking into consideration that the sample with both proteins contained a significantly larger number of droplets and no visible aggregates, there existed a number of possible explanations. First, the cross-interaction between both proteins could stabilize S100A9 and prevent its aggregation, which would explain the lack of amorphous structures and a higher number of droplets. Second, the S100A9 aggregates may be present in the sample, but they are not visible due to their inability to interact with eGFP-α-syn. Lastly, S100A9 may associate with α-syn into droplets and then rapidly form aggregates, which would explain the previously observed uneven distribution in part of the condensates (Figure 1E).

To answer this question, S100A9 was aggregated into fibrils prior to being combined with α-syn and eGFP-α-syn. During sample analysis with brightfield microscopy, the first notable observation was that S100A9 fibrils, which are normally short worm-like structures ^50^, associated into large amorphous aggregate clusters (Figure 4A). These S100A9 aggregate assemblies ranged from several to well over a hundred micrometers in size. Another interesting factor was that the S100A9 aggregates were clearly visible during fluorescence microscopy due to their association with eGFP-α-syn (Figure 4B). These results indicate that the hypothesis of S100A9 aggregation outside of α-syn droplets is not correct, as they would be clearly visible in the fluorescence microscopy images. The possibility of rapid S100A9 aggregation within heterotypic droplets remains inconclusive, however, the observed large size of S100A9 structures under high molecular crowding conditions also makes this event unlikely. Additionally, the fluorescence microscopy images revealed that there is a high level of cross-interaction between α-syn and S100A9 aggregates.

**Figure 4.**
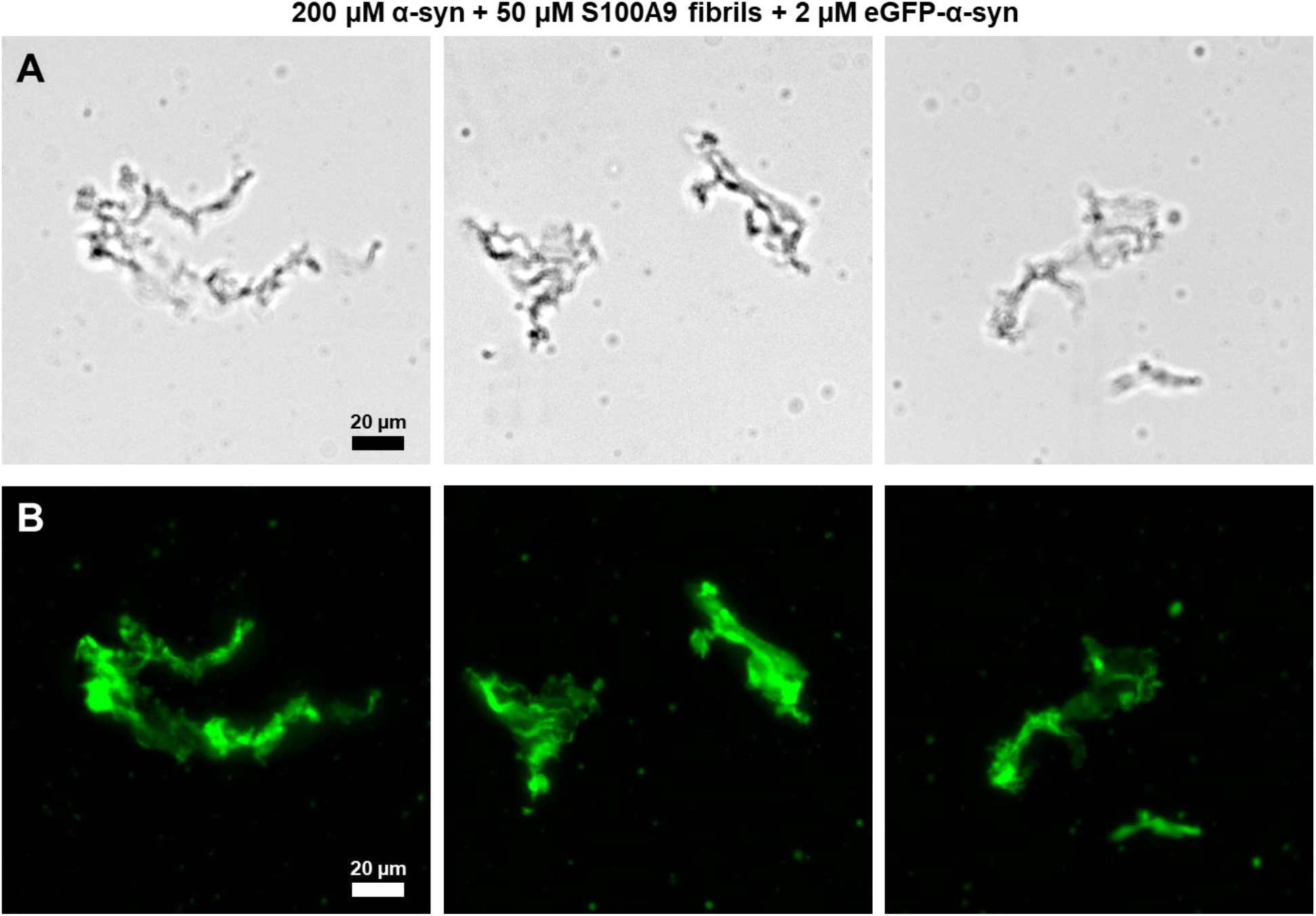
Brightfield and fluorescence microscopy images of S100A9 aggregates with α-syn. Brightfield microscopy images (Olympus IX83 microscope) of samples containing 200 µM α-syn, 50 µM S100A9 fibrils and 2 µM eGFP-α-syn (A, scale bar – 20 µm). Fluorescence microscopy images of the samples at the same exact positions (B, scale bar – 20 µm).

Since the results of this study indicated that both native, as well as aggregated S100A9 can interact with α-syn under LLPS-inducing conditions, it was further investigated whether this cross-interaction can influence the process of α-syn fibril formation. Samples containing different concentrations of S100A9 and its aggregated form with or without α-syn were monitored under LLPS-inducing conditions with the use of an amyloid-specific dye – thioflavin-T (ThT). In the case of the S100A9 control samples, the native protein either displayed a very low increase in ThT fluorescence intensity (at 5 µM concentration, Figure 5A, C) or had a rapid initial increase, followed by a second change in fluorescence intensity (at 50 µM concentration, Figure 5A, C). The double transition in ThT fluorescence intensity can be attributed to the initial assembly of droplets/aggregates, after which amyloid fibrils are formed ^44^. Interestingly, when the S100A9 aggregates (prepared under non-LLPS conditions) were subjected to the same conditions (Figure 5B), the 50 µM sample end-point fluorescence intensity values were only half of what was observed in the case of the native protein sample (Figure 5C). This result indicates that LLPS conditions may induce the formation of a different S100A9 aggregate conformation with distinct ThT-binding properties ^54^.

**Figure 5.**
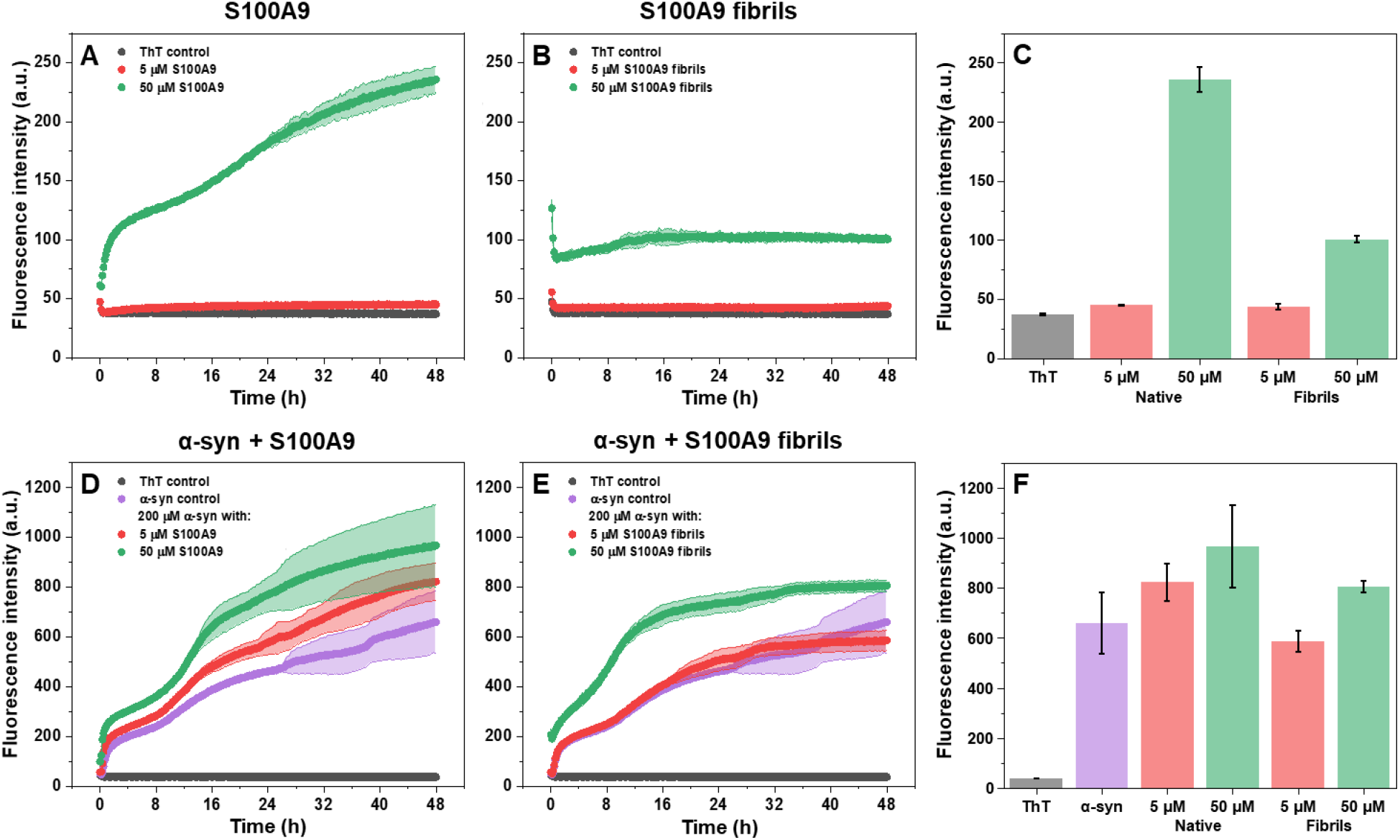
LLPS and aggregation kinetics of α-syn with S100A9. Native S100A9 (A), S100A9 fibril (B), α-syn with native S100A9 (D) and α-syn with S100A9 fibrils (E) sample ThT fluorescence intensity changes over 48 hours of incubation under LLPS-inducing conditions. End-point fluorescence intensity values of samples after 48 hours of incubation (C, F). Error plots and bars are for one standard deviation (4 technical repeats for each condition).

When α-syn was combined with 5 or 50 µM of native S100A9, the ThT intensity kinetic curves followed a similar double-sigmoidal trend, with the only exception being the fluorescence intensity values (Figure 5D). In the case of S100A9 fibrils, the kinetic curves of the α-syn control and 5 µM aggregated S100A9 were nearly identical (Figure 5E). Surprisingly, the second increase in ThT fluorescence intensity occurred much quicker when the sample contained 50 µM S100A9 fibrils, which may be related to aggregate surface-enhanced secondary nucleation ^55^. This sample also yielded a higher end-point fluorescence intensity value, similar to the one with native S100A9 (Figure 5F). The differences between these end-point intensity values can either stem from the formation of structurally distinct α-syn fibrils, as was previously shown with this protein pairing ^41^, or they can be the result of S100A9 aggregate formation. In order to figure this out, the α-syn samples were subjected to two rounds of reseeding. As self-replication is a hallmark of amyloid structures, the two rounds of reseeding (10% seed) would simultaneously increase the abundance of α-syn fibrils, greatly diminish the number of amorphous aggregates, as well as massively reduce the concentration of S100A9 fibrils and PEG in the samples.

During the first round of reseeding, α-syn fibrils (initially prepared with 5 or 50 µM native S100A9) both displayed exponential aggregation curves (Figure 6A), which were similar in shape to the control and indicated efficient reseeding. Both samples had a much larger end-point fluorescence intensity than the control (Figure 6C), which supported the hypothesis that native S100A9 can lead to the formation of distinct α-syn fibril structures. Oppositely, α-syn aggregates formed in the presence of S100A9 fibrils produced a lower end-point fluorescence intensity than the control (Figure 6B, C), suggesting that either the protein structures had a different ThT-binding mode or that the self-replication was highly inefficient.

**Figure 6.**
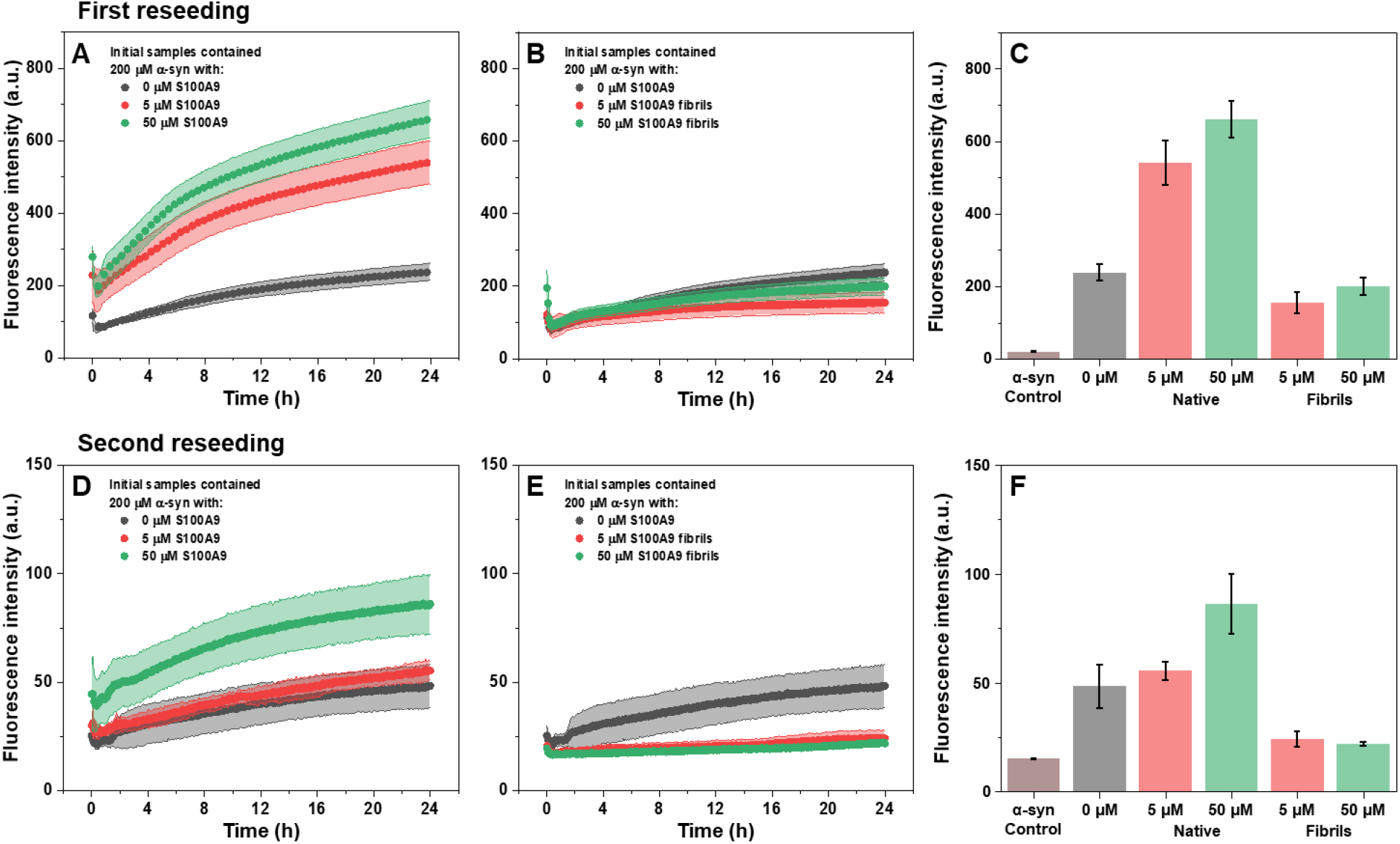
α-syn aggregate reseeding kinetics and end-point fluorescence intensity values. First and second round of α-syn aggregate reseeding after their preparation in the presence of native (A, D) or aggregated (B, E) S100A9. End-point fluorescence intensity values of the first (C) and second (F) round of reseeding (monomeric α-syn sample intensity added for comparison). Error plots and bars are for one standard deviation (6 technical repeats for each condition). Larger scale panel E kinetics are available as Figure S4.

The second round of reseeding resulted in an even higher disparity between the results. In the case of α-syn aggregates prepared in the presence of native S100A9, only the sample which initially contained 50 µM S100A9 resulted in a notable increase in ThT fluorescence intensity, while the 5 µM initial S100A9 sample was similar to the control (Figure 6D, F). As in the first reseeding, the ThT fluorescence intensity increase of samples initially prepared with aggregated S100A9 followed a similar trend (Figure 6E, F). The sample signal intensity was only slightly higher than the α-syn monomer control and much lower than the α-syn aggregate control (Figure S4). These results indicated that α-syn aggregates were only capable of efficient self-replication when they were initially prepared in the absence of S100A9 or in combination with native S100A9.

To evaluate what structural differences occurred due to the formation of heterotypic α-syn and S100A9 droplets, the α-syn control and α-syn with 50 µM S100A9 samples (after two rounds of reseeding) were selected for further investigation by Cryo-EM. Out of the three, the fibril content of the sample initially prepared with S100A9 aggregates was insufficient for an accurate determination of their structures. In the case of the control sample, it contained two structurally distinct fibrils with the dominant type constituting approximately 92% of the total distribution (Figure 7A, S5). When the initial α-syn fibrils were prepared in the presence of 50 µM S100A9, the entire reseeded sample consisted of fibrils with an identical morphology (Figure 7B, S6). This dominant fibril class was the same under both conditions, indicating that the presence of native S100A9 can reduce the variability of α-syn aggregate formation.

**Figure 7.**
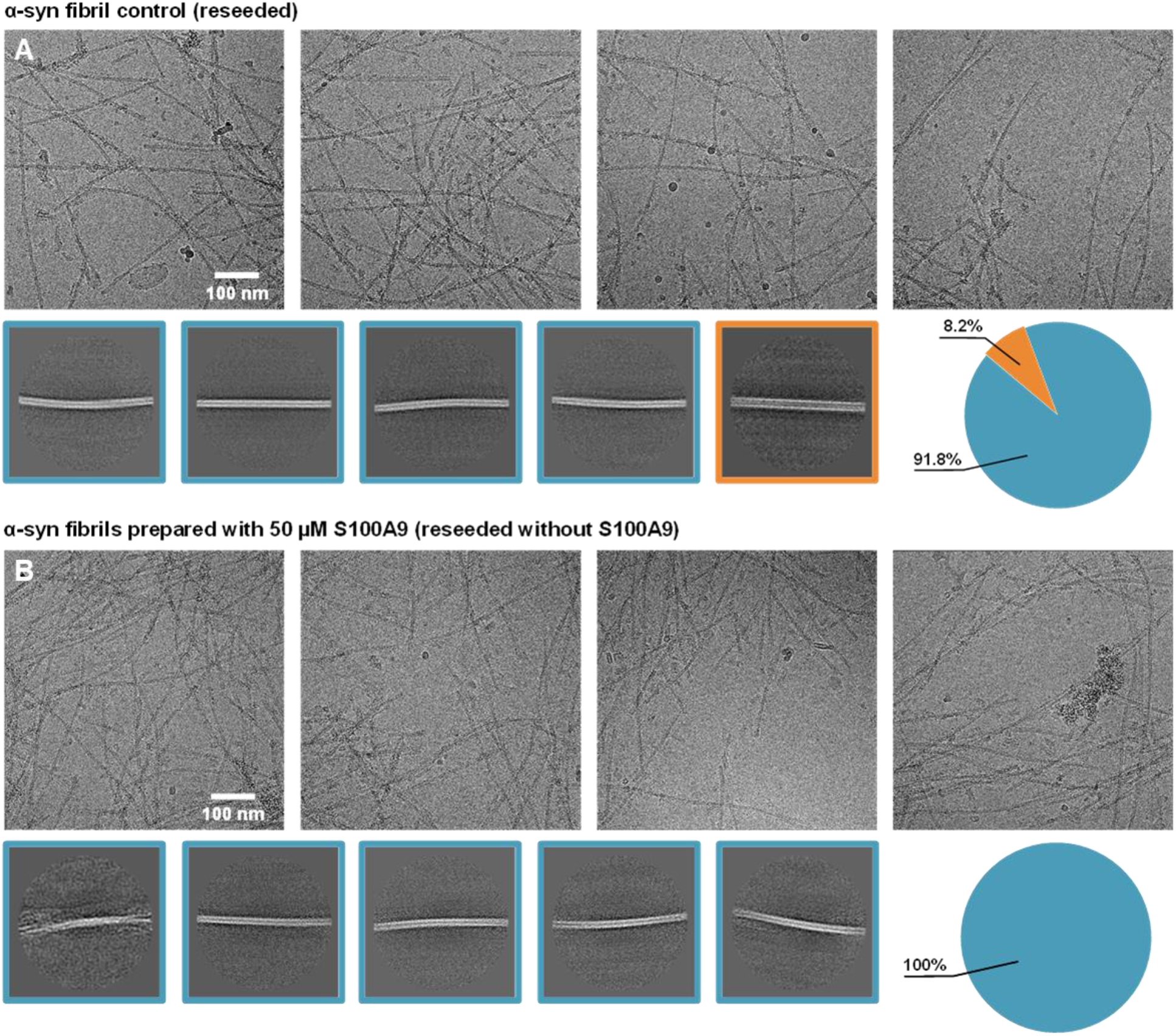
Cryo-EM images of α-syn fibrils after two rounds of reseeding. Images of α-syn fibrils which were prepared in the absence (A) or presence (B) of 50 µM native S100A9 and then reseeded twice (10% seed (v/v) each time). Different fibril classes are indicated by blue and orange outlines. Fibril distributions are shown as color-coded pie-charts next to the images.

## Discussion

The cross-interaction between S100A9 and α-syn has previously been observed to modulate the amyloid formation process of α-syn in vitro ^41^, as well as their tendency to co-localize in aggregate plaques in vivo ^32^. In this work, we examined whether this interaction could also lead to the formation of heterotypic protein condensates, a phenomenon previously observed for certain sets of amyloidogenic proteins. We discovered that α-syn and S100A9 can associate into condensates under high molecular crowding conditions and observed several interesting aspects regarding their liquid-liquid phase separation.

Previous reports of amyloid protein heterotypic droplet formation have generally shown a homogenous distribution of both molecules within condensates ^17,19,20^. This did not appear to be the case in respect to α-syn and S100A9, where we observed the formation of regular homotypic droplets alongside their heterotypic variants. Fluorescence microscopy images of α-syn samples containing mCherry-S100A9 also displayed an uneven distribution of the labeled protein within the condensate. These findings would suggest that both proteins can exist within the same droplets, however, they may have a higher level of affinity towards self-association.

Additionally, a number of interesting observations were made in the case of S100A9 within a high molecular crowding environment. Both the fluorescently-labeled and regular S100A9 displayed a very high level of self-association under LLPS-inducing conditions, including the formation of droplets and aggregates. We have previously observed that fluorescently-labeled proteins can possess a higher level of condensate formation ^46^, however, this effect appeared to be exceptionally high for mCherry-labeled S100A9. The high molecular crowding environment also had an interesting influence on S100A9 aggregates. While they typically exist as short worm-like structures, the LLPS-inducing conditions caused them to clump into large aggregate structures. These assemblies were highly efficient at binding α-syn and even enhancing their transition to amyloid fibrils via surface-mediated nucleation.

Another interesting aspect of this cross-interaction was the stabilization of a single dominant α-syn fibril strain. Such an effect of S100A9 has previously been observed in vitro, where the mixture of both proteins resulted in the generation of a single α-syn aggregate secondary structure ^41^. Taking into consideration that only a part of S100A9 and α-syn formed heterotypic droplets, it is likely that this influence on the resulting fibril structure is not entirely dependent on the protein association within droplets. Since the relative abundance of α-syn aggregation centers is low during the process of unseeded nucleation, the presence of even a low concentration of S100A9 within or outside of the condensates may influence nuclei formation and elongation.

The reseeding experiments also revealed an unexpected result of α-syn and S100A9 heterotypic droplet formation. When both proteins were in their native state within the condensate, α-syn aggregated into fibrils which pertained a notably higher level of self-replication properties when subjected to a non-LLPS-inducing environment. Taking into consideration the localization of both proteins, as well as the high molecular crowding environment in vivo, it is possible that heterotypic droplet formation results in fibrils with a higher tendency for self-replication. This would suggest that heterotypic droplet formation between the two proteins may be a critical step in the onset and progression of α-syn-related neurodegenerative disorders.

## Conclusions

The pro-inflammatory S100A9 and neurodegenerative disease-related alpha-synuclein are capable of forming heterotypic droplets under LLPS-inducing conditions. This cross-interaction can lead to a stabilization of a specific α-syn fibril strain, which is capable of effective self-replication under non-LLPS conditions. The ability of these proteins to form heterotypic condensates complements the previously reported cross-interactions in vivo and in vitro, adding another piece to the complex amyloid interactome puzzle.

## Data availability

All raw data and additional images are available in an online data repository at: https://data.mendeley.com/datasets/tvf9nwtdhn/1.

## Author contributions

Conceptualization – M.Z., D.S., V.S.; Investigation – D.V., A.K., D.S., M.Z., K.M.; Data curation – M.Z.; Writing – Original draft preparation – M.Z.; Writing – review & editing – D.V., A.K., D.S., K.M., M.T., V.S., M.Z.; Visualization – M.Z., D.S., A.K.; Supervision – M.Z., M.T., V.S.

