## Supporting information for "Heterotypic droplet formation by pro-inflammatory S100A9 and neurodegenerative disease-related alpha-synuclein"

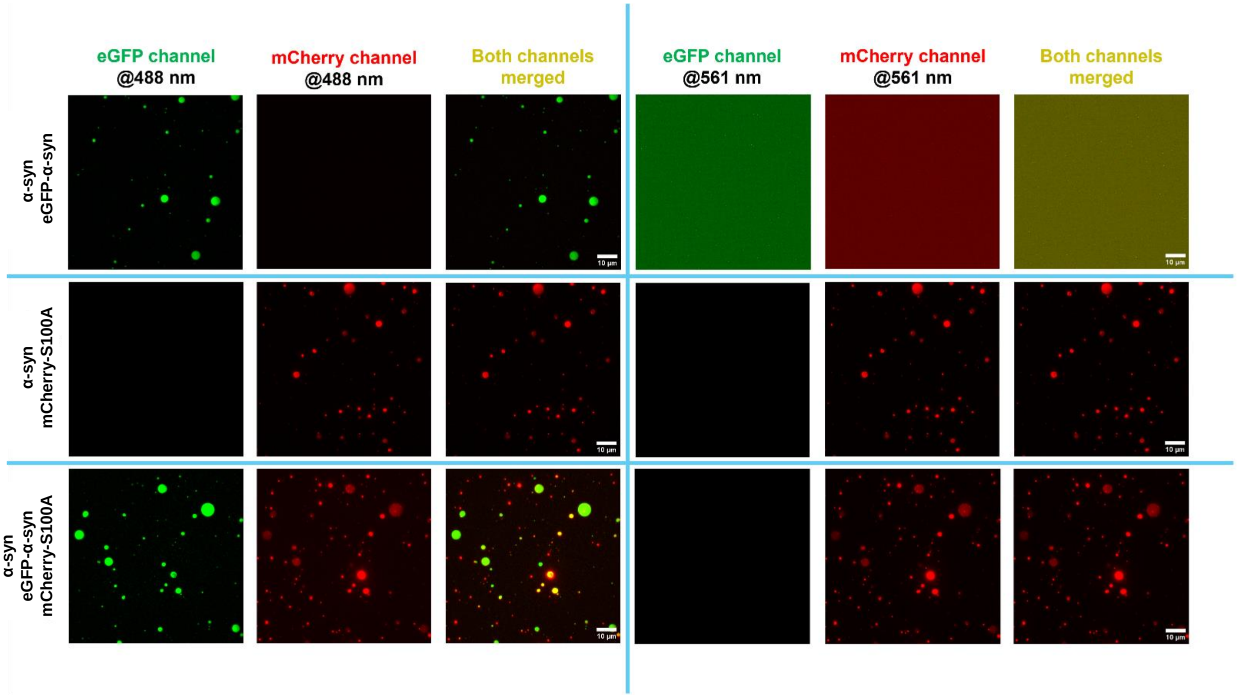


**Figure S1.** TIRF microscopy images of droplets formed from α-syn with either eGFP-α-syn, mCherry-S100A9 or both. Such droplets were immobilized on the glass surface and visualized in the eGFP and mCherry spectral channels using either 488 nm or 561 nm laser excitation. For each fusion construct, the same imaged surface position is shown and every set of images separated by the blue lines have identical minimum and maximum intensity scale values applied to them.

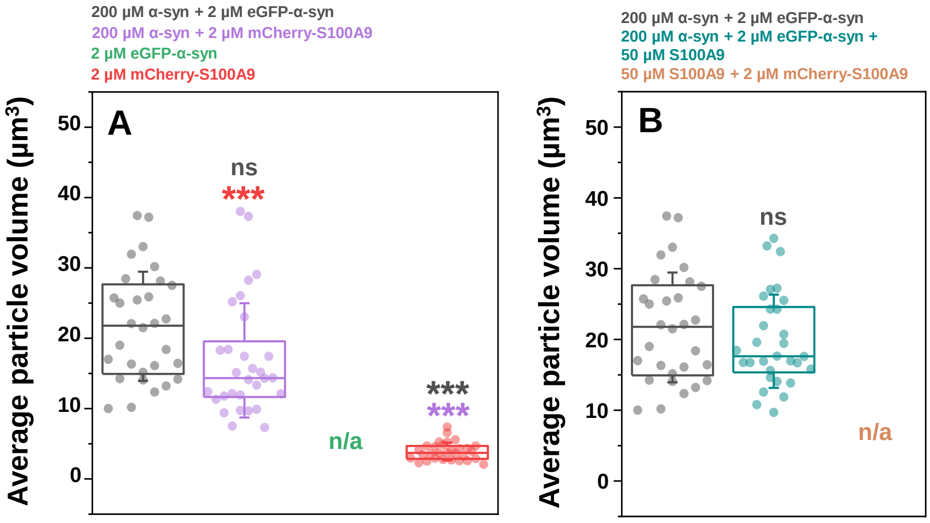


**Figure S2.** Statistical analysis (ANOVA Bonferroni means comparison, thirty 500x500 pixel size images, ns – not significant, *** $p<0.001$) of the particle average volume distribution in the absence (A) or presence (B) of 50 μM native S100A9.

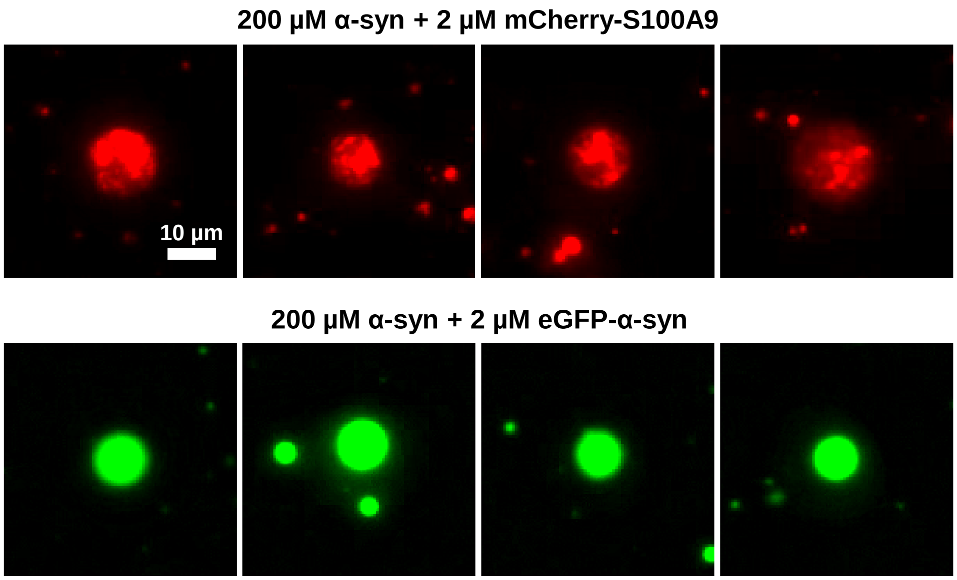


**Figure S3.** Comparison of labeled protein distribution within α-syn droplets. Scale bar (10 μm) is identical for all images.

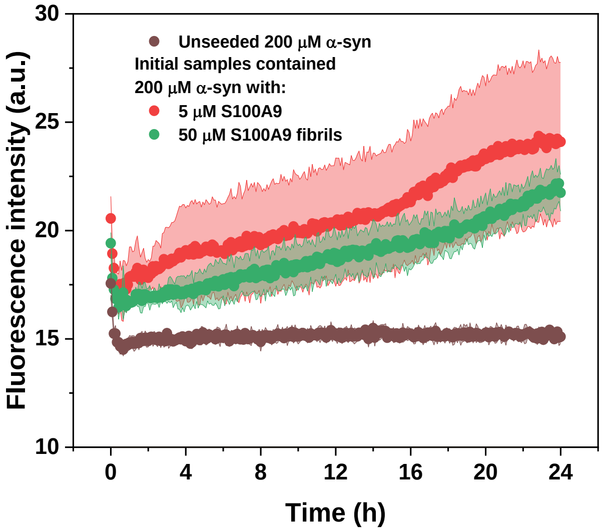


**Figure S4.** Second round of α-syn aggregate reseeding kinetics.

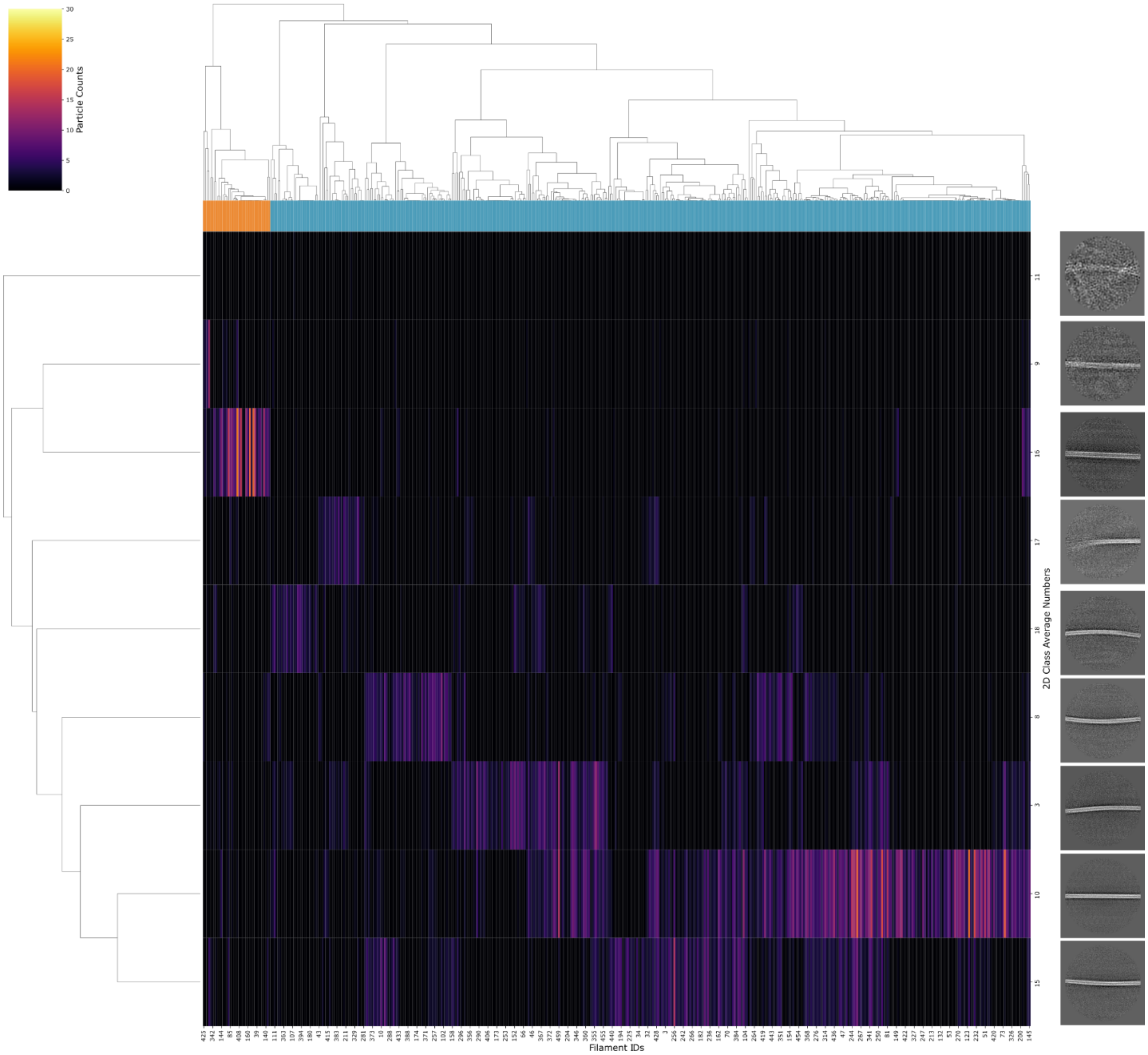


**Figure S5.** Hierarchical classification of α-syn (reseeded from α-syn LLPS) filament segments according to their assigned 2D class average number (vertical) and the picked filament ID (horizontal).

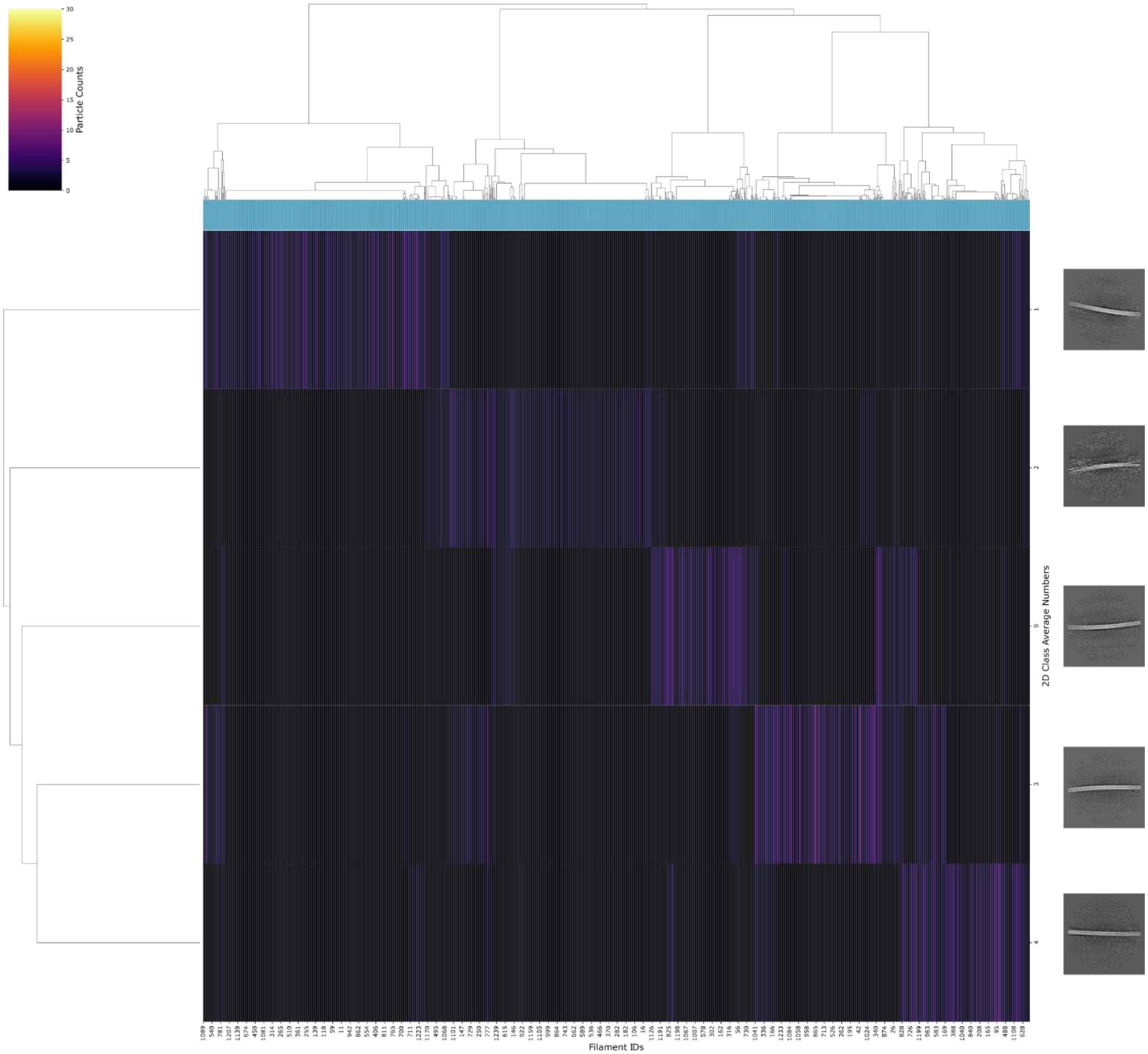


**Figure S6.** Hierarchical classification of α-syn (reseeded from α-syn + S100A9 LLPS) filament segments according to their assigned 2D class average number (vertical) and the picked filament ID (horizontal).

| Primers | Sequence |
|---|---|
| DS105_S100A9_frw | GTGGTGGTGGTTCTGGTGGTGGTGGTTCTATGACTTGCAAAATGTCGCAGCTGG |
| DS105_S100A9_rev | GCGGATCCTTAGGGGGTGCCCTCCCCG |
| DS105_mCherry_frw | CGCATATGGTGAGCAAGGGCGAAGAAGATAAC |
| DS105_mCherry_rev | GAACCACCACCACCAGAACCACCACCACCCTTGTACAGCTCGTCCATGCCGCCG |

**Table S1**. Primers used in this study

| Name | α-syn (reseeded from α-syn LLPS) | α-syn (reseeded from α-syn +S100A9 LLPS) |
|---|---|---|
| **Data Collection** | | |
| Pixel Size (Å) | 1.1 | 1.1 |
| Defocus range (nm) | -2.2 to –1.2 | -2.2 to –1.2 |
| Voltage (kV) | 200 | 200 |
| Camera | Falcon 3EC | Falcon 3EC |
| Microscope | Glacios | Glacios |
| Exposure time (s) | 46.33 | 46.33 |
| Number of frames | 30 | 30 |
| Total dose ($e^-$/Å$^2$) | 30 | 30 |
| **2D Classification** | | |
| Micrographs | 881 | 973 |
| Picked fibrils | 1158 | 8816 |
| Box size (px) | 1024 | 1024 |
| Inter-box distance (Å) | 69.09 | 69.09 |
| Segments Extracted | 4072 | 9204 |
| Segments used for final 2D classification | 4072 | 4060 |

**Table S2.** Statistics of Cryo-EM data collection and 2D classification.
